# A UNIVERSAL FRAMEWORK FOR DISENTANGLING SUBJECT-SPECIFIC SIGNATURES IN EEG SIGNALS

**DOI:** 10.64898/2026.05.26.727876

**Authors:** Zhaodi Pei, Ziyu Li, Qing Li, Xia Wu

## Abstract

Extracting stable subject-specific features from EEG signals remains challenging due to their entanglement with transient brain states. We propose a universal neural framework that disentangles subject-specific features from state-dependent components in raw EEG signals. Our approach employs a disentanglement module with a cross-reconstruction objective to isolate subject-specific representations. We validate our framework on EEG-based biometric recognition using two public datasets with leave-one-state-out cross-validation. Results demonstrate significant improvements in out-of-distribution identification accuracy across four different backbone models, confirming our method’s universality and plug-and-play capability. This work advances reliable extraction of neural signatures for personalized neurotechnology applications.

## 1. INTRODUCTION

Extracting stable, subject-specific signatures from brain signals is pivotal for personalized neurotechnology [1, 2]. These neural “brainprints” enable personalized diagnostics, rapid BCI calibration, and robust biometrics [3, 4, 5]. Among acquisition methods, Electroencephalography (EEG) is ideal for real-world deployment due to its non-invasiveness and temporal precision [6].

Both traditional engineered features [7] and deep learning models [8] often entangle identity information with transient neural activity, degrading performance on out-of-distribution (OOD) states [9]. While prior state-independent methods attempted to mitigate this via assumption-driven feature engineering (e.g., subspace projections) [10], they lack the flexibility of end-to-end learning, highlighting the need for automatic extraction of state-invariant features.

To address this generalization barrier, we propose a universal framework to disentangle subject-specific signatures from state-dependent features. Our approach is grounded in the causal assumption that subject identity represents an invariant factor while brain states vary. The core is a novel disentanglement module using a cross-reconstruction objective: reconstructing a subject’s EEG in one state using subject-specific features from a different state. This forces the learning of state-invariant representations. We further propose a discrimination module to enhance inter-subject differences. Validated on OOD biometric recognition via leave-one-state-out cross-validation (LOSO-CV), our method achieves state-of-the-art performance in two multiple states datasets. Crucially, it functions as a “plug-and-play” module, significantly improving OOD generalization across four distinct backbone models, offering a robust solution for reliable EEG applications.

## 2. METHOD

### 2.1. Problem Formulation

We address the challenge of disentangling stable subject-specific neural signatures from transient state-dependent brain activity in EEG signals. Each EEG segment **X** ∈ℝ^*C×T*^ (*C* channels, *T* temporal length) from subject *k* during cognitive state *p* comprises two latent factors: a subject-specific signature **z**_**id**_ and a state-dependent component **z**_**non**_**id**_.

Our objective is to learn encoder networks that disentangle these factors:

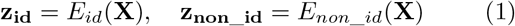

This disentanglement must satisfy: (1) State Invariance: *E*_*id*_(**X**^*k,p*^) *≈ E*_*id*_(**X**^*k,q*^) for the same subject *k* under different states; (2) Subject Discriminability: *E*_*id*_(**X**^*k*^) ≠ *E*_*id*_(**X**^*h*^) for different subjects; and (3) Information Preservation: (**z**_**id**_, **z**_**non**_**id**_) collectively preserve information to reconstruct **X**.

### 2.2. Disentanglement Module

We achieve disentanglement through cross-reconstruction using two specialized encoders, *E*_*id*_ and *E*_*non*_*id*_, and a shared decoder *D*. While both encoders share identical architectures, they maintain distinct parameters to capture complementary information: *E*_*id*_ extracts subject-specific representation, and *E*_*non*_*id*_ captures state-dependent representation.

The cross-reconstruction mechanism operates on paired EEG signals from the same subject under different cognitive states: *X*^*k,p*^ and *X*^*k,q*^ (subject *k* in states *p* and *q*, respectively). For more than two states, we employ an “all-pairs” strategy. Each signal is encoded into two latent representations:

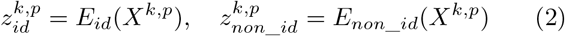

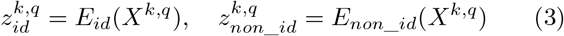

The key insight is that subject-specific attributes should remain consistent across states, while state-dependent attributes should vary. We therefore reconstruct each signal by combining its own state-dependent component with the subject-specific component from the paired signal:

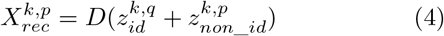

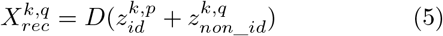

This cross-reconstruction is enforced through the reconstruction loss:

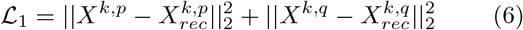

However, solely minimizing ℒ_1_ may lead to a trivial solution where *E*_*non*_*id*_ reconstructs the entire signal while *E*_*id*_ collapses. To prevent this and ensure meaningful subject representations, we impose additional constraints to enforce discriminative identity features.

### 2.3. Discrimination Module

To ensure that subject-specific attributes contain discriminative identity signatures, we introduce a discrimination module consisting of a two-layer fully connected classifier that maps *z*_*id*_ to identity labels. This module also addresses the potential shortcut learning issue in the disentanglement module by enforcing explicit intra-subject discrimination.

The discrimination loss operates on both original and reconstructed subject-specific representations. Since successful cross-reconstruction should preserve identity information, we require that reconstructed subject-specific attributes 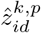and 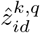 maintain the same identity as their original counterparts. The discrimination loss is formulated as:

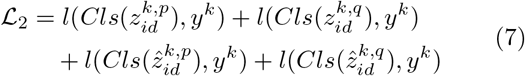

where *Cls*(·) denotes the identity classifier, *l*(*·*) represents cross-entropy loss, and *y*^*k*^ is the identity label for subject *k*, and identity labels are encoded as one-hot vectors.

The two modules work synergistically: the disentanglement module separates subject-specific from state-dependent factors to achieve task-independence, while the discrimination module ensures subject-specific attributes remain discriminative for identity recognition. Together, they enable robust state-invariant EEG-based biometric identification.

### 2.4. Implementation of Uni-SEB

The overall loss function of Uni-SEB combines disentanglement and discrimination objectives:

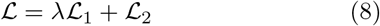

where *λ* = 1 balances both objectives. We employ paired sampling (same subject, different states) with 32 pairs per batch to facilitate feature disentanglement. The model is trained for 500 epochs using Adam optimizer (learning rate=1 × 10^*−*3^), with the final model selected based on peak validation accuracy.

## 3. EXPERIMENTS AND RESULTS

### 3.1. Datasets and Preprocessing

We evaluate Uni-SEB on two multi-state datasets: the public Biometric EEG Dataset (BED) [11] and our self-collected MSEEG^1^. BED includes 14-channel EEG from 21 subjects across 6 states (e.g., resting, VEP) recorded at 256 Hz over three sessions. MSEEG features 32-channel data from 28 subjects across 3 tasks, including motor imagery, working memory, and cognitive load tasks.

For preprocessing, we applied the PREP pipeline [12] with downsampling to 128 Hz for BED, and FIR filtering (0.3–45 Hz) with downsampling from 500 Hz to 100 Hz for MSEEG. Downsampling is employed for computational efficiency, adhering to standard practices in EEG deep learning. Following [7], all signals were segmented into non-overlapping 1s windows.

### 3.2. Experimental Setup

To assess the stability of neural signatures across cognitive contexts, we employ a LOSO-CV protocol. In each fold, one state serves as the OOD test condition, while the remaining states are used for training (split 7:1:2 for train/val/test). Performance is measured by the average identification accuracy (proportion of correctly identified samples) on the unseen OOD states across all iterations.

### 3.3. Comparison methods

We benchmark Uni-SEB against traditional features— such as Power Spectral Density (PSD), Mel-Frequency Cepstral Coefficients (MFCCs), and Autoregressive (AR) coefficients—using SVM classifiers, as well as deep networks, including LSTM [13], GRU [14], EEGNet [15], TSCeption [16] and EEGNeX [17]. Note that Multilayer Perceptrons (MLPs) were excluded due to their excessive computational cost. To demonstrate Uni-SEB’s architecture-agnostic utility, we integrate it into the four aforementioned backbones, evaluating its ability to boost generalization without structural modifications. For the RNN models (LSTM and GRU), we employ a single-layer encoder with a tanh activation and a dropout rate of 0.5. For EEGNet and TSCeption, we adhere to the configurations specified in their original papers.

### 3.4. Generalization performance in OOD cognitive states

We evaluate Uni-SEB under the LOSO-CV protocol on both datasets (Table 1). While traditional features (AR, MFCC, PSD) struggle with generalization, integrating Uni-SEB consistently boosts end-to-end baselines. Specifically, Uni-SEB+TSCeption achieves the highest accuracy on the BED dataset (91.37%), and Uni-SEB+EEGNet leads on the MSEEG dataset (97.17%). Notable gains are also observed in RNNs on both datasets. These results across diverse backbones confirm Uni-SEB’s universality in extracting state-independent signatures.

**Table 1.** The overall OOD identification accuracy of different methods with the LOSO-CV experimental setup.

| Methods |  | Overall OOD Identification Accuracy |  |
| --- | --- | --- | --- |
|  |  | BED Dataset | MSEEG Dataset |
| Hand-Crafted Features | AR | 59.31 ± 2.34% | 85.11 ± 3.42% |
|  | MFCC | 57.95 ± 2.39% | 60.77 ± 2.34% |
|  | PSD | 48.21 ± 4.97% | 84.04 ± 2.57% |
| End-to-End Methods by RNNs | LSTM | 66.38 ± 5.40% | 87.77 ± 5.75% |
|  | Uni-SEB+LSTM | 78.28 ± 5.58% | 91.04 ± 4.91% |
|  | GRU | 75.47 ± 3.19% | 83.30 ± 2.30% |
|  | Uni-SEB+GRU | 83.17 ± 3.54% | 88.60 ± 2.49% |
| End-to-End Methods by CNNs | EEGNet | 87.99 ± 2.10% | 96.25 ± 1.96% |
|  | Uni-SEB+EEGNet | 91.05 ± 1.57% | 97.17 ± 2.17% |
|  | TSception | 88.12 ± 2.01% | 86.37 ± 4.57% |
|  | Uni-SEB+TSception | 91.37 ± 1.92% | 90.83 ± 4.06% |
|  | EEGNeX | 78.41 ± 3.38% | 82.97 ± 3.58% |
|  | Uni-SEB+EEGNeX | 79.18 ± 2.57% | 86.63 ± 3.16% |

<sup>1</sup>Approved by Southwest University Ethics Committee.

### 3.5. Visualization of spatial features for brainprint identification with Uni-SEB

We employ saliency-based topological maps to interpret spatial contributions to brainprints. Taking Subject 01 on both datasets for example, Fig. 2 (BED) and Fig. 3 (MSEEG) consistently highlight the occipital (O2) and frontal (F4 for BED, FC1 for MSEEG) lobes as critical regions across cognitive states. The stability of these areas—associated with visual processing and decision-making—validates Uni-SEB’s robustness in isolating state-independent identity signatures.

**Fig. 1.**
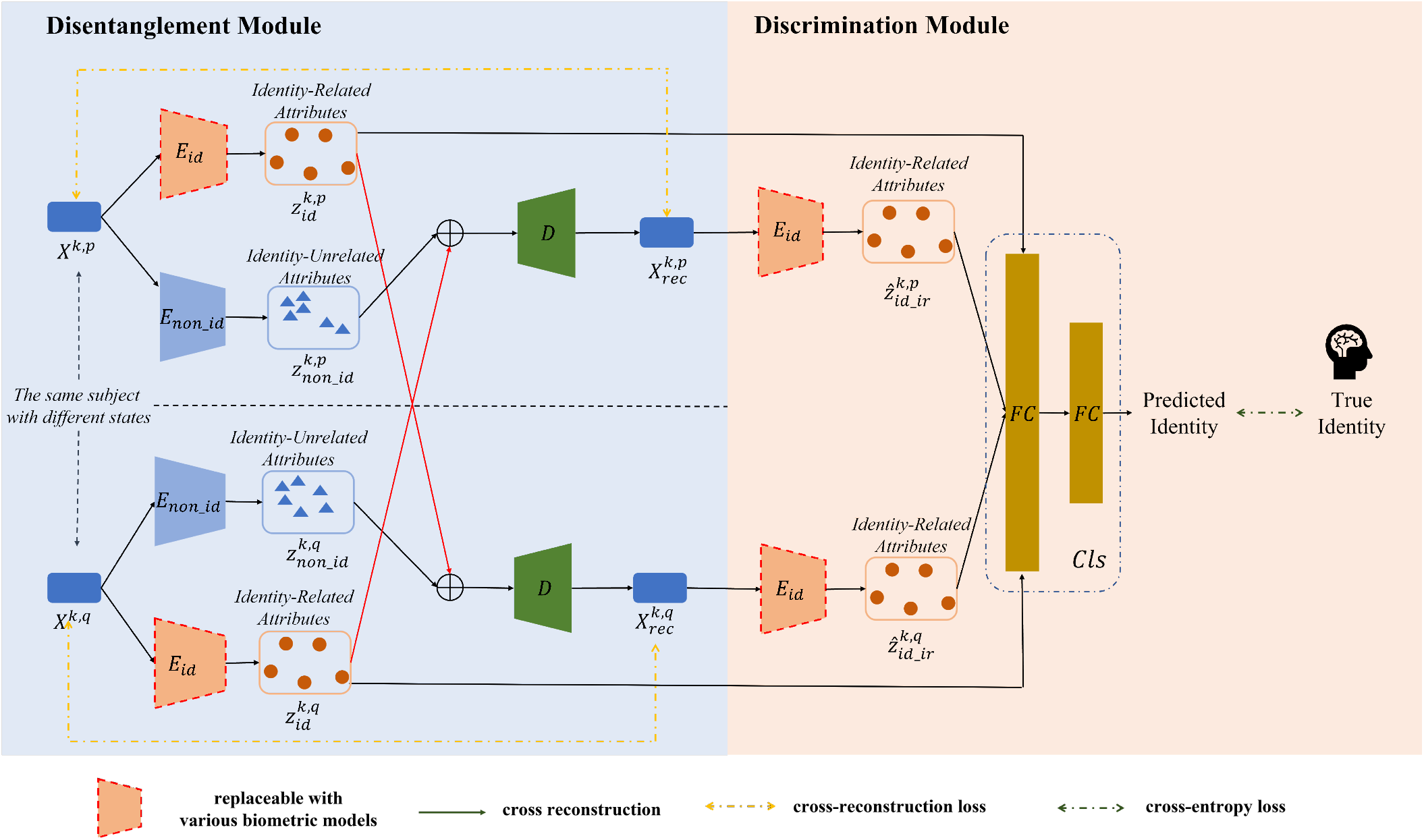
Overview of the proposed Uni-SEB training procedure. The disentanglement module enforces consistency of subject-specific features across states via cross-reconstruction, while the discrimination module enhances inter-subject separability through individual identification.

**Fig. 2.**
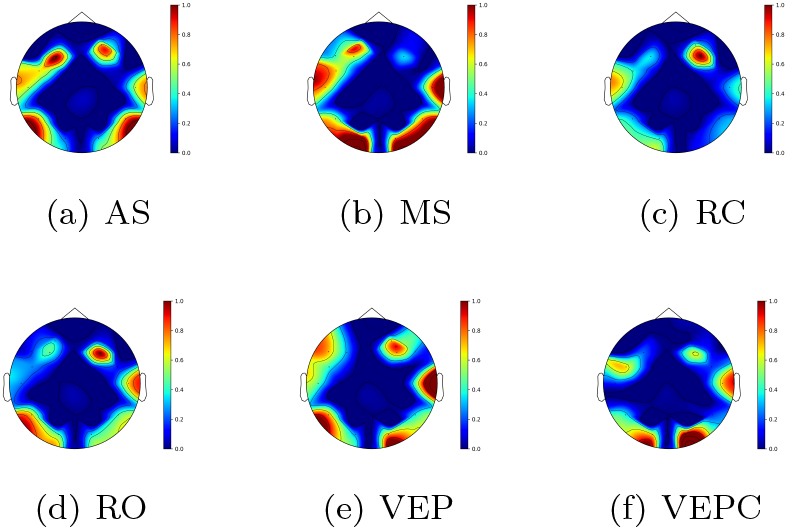
Spatial contribution topomaps for Subject 01 across six states in BED.

**Fig. 3.**
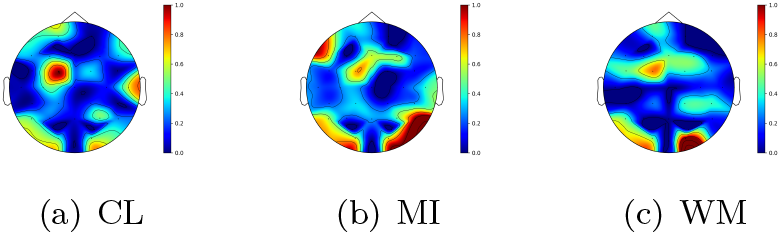
Spatial contribution topomaps for Subject 01 across three states in MSEEG.

### 3.6. Hyperparameter Sensitivity Analysis

We investigate the impact of the *λ* on Uni-SEB performance across values {0.2, 0.5, 1, 2, 5}.As shown in Fig. 4, both datasets exhibit consistent patterns with optimal accuracy achieved at *λ* = 1. Performance declines slightly as *λ* deviates from this value, confirming that *λ* = 1 strikes the optimal balance between feature disentanglement and identification objectives.

**Fig. 4.**
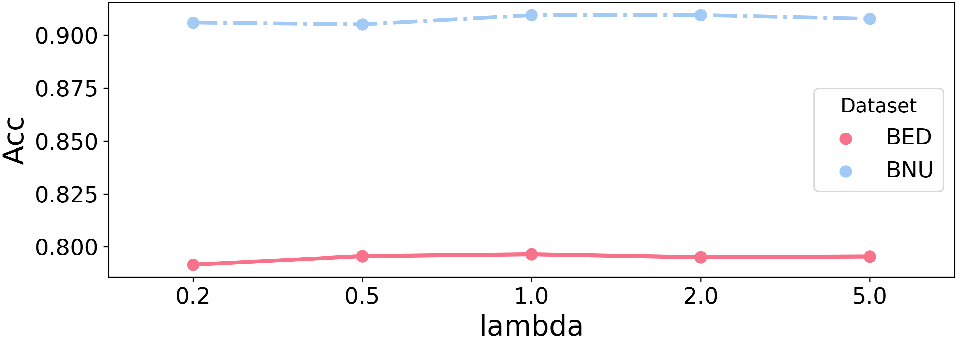
The influence of the hyperparameter *λ*.

### 3.7. Ablation Study

We analyze component contributions using Uni-SEB+LSTM (Table 2). The w/o Disent variant performs similarly to the LSTM baseline. This is expected, as removing the disentanglement module structurally reduces the framework to a basic encoder, confirming that our generalization gains stem directly from the proposed disentanglement mechanism. Conversely, w/o Discrim causes poor accuracy; while it captures intra-subject consistency, it lacks the inter-subject supervision necessary for feature separability. These results validate the necessity of our joint optimization strategy.

**Table 2.** Ablation study results.

|  | BED | MSEEG |
| --- | --- | --- |
| w/o Discrim | 4.13±0.58% | 29.38±9.11% |
| w/o Disent | 66.38 ± 5.40% | 87.77±5.75% |
| Uni-SEB | 78.28 ± 5.58% | 91.04±4.91% |

## 4. CONCLUSION

We proposed a universal framework that disentangles subject-specific signatures from EEG signals via a novel cross-reconstruction module. By maintaining feature consistency across cognitive states, our approach effectively isolates stable “brainprints” independent of mental tasks. While this study prioritizes the challenge of task-independent variability, it establishes a solid foundation for future cross-session research. Ultimately, this work advances the robust isolation of individual neural signatures, facilitating reliable personalized neuroscience.

## 5. ACKNOWLEDGEMENT

This work is supported by the National Science Fund for Distinguished Young Scholars (Grant Number 62325601).

## Footnotes

1 Approved by Southwest University Ethics Committee.

